# Potential of breadfruit cultivation to contribute to climate-resilient low latitude food systems

**DOI:** 10.1101/2021.10.01.462801

**Authors:** Lucy Yang, Nyree Zerega, Anastasia Montgomery, Daniel E. Horton

## Abstract

The number of people in food crisis around the world is increasing, exacerbated by the challenges of COVID-19 and a rapidly changing climate. Major crop yields are projected to decrease in low-latitude regions due to anthropogenic climate change, making tropical and sub-tropical food systems particularly vulnerable to climate shocks. Increased cultivation of breadfruit (*Artocarpus altilis*), often categorized as a neglected and underutilized species (NUS), has been suggested as an agricultural adaptation pathway for food insecure tropical and subtropical regions, due to its potential to enhance climate resilience and overall sustainability of low-latitude agricultural systems. To better understand breadfruit’s cultivation suitability and geographic range in current and future climates, we employ a diverse set of observations and models to delineate the current climatically viable breadfruit range and assess the climatically viable breadfruit range in the future (2061-2080) under stabilization and high emission scenarios. We find that the area of suitable breadfruit range within the tropics and subtropics is projected to decrease ~4.4% in the stabilization scenario and ~4.5% in the high emission scenario. In Southeast Asia and the Pacific Islands, yield quality and consistency show minimal decreases under the high emission scenario, with increases in total suitable area under both scenarios. In contrast, in Latin America and the Caribbean, the current range of breadfruit suitability is projected to contract ~10.1-11.5% (stabilization-high emission). Present and future model suitability outputs suggest that opportunities to successfully expand breadfruit cultivation over the next several decades exist in sub-Saharan Africa, where food insecurity is coincidentally high. However, in all regions, high emission scenario conditions reduce the overall consistency and quality of breadfruit yields compared to the stabilization scenario. Our results have the potential to inform global food security adaptation planning and highlight breadfruit as an ideal NUS to incorporate in food security adaptation strategies in a changing climate.

## 1 Introduction

The number of people in the world lacking regular access to nutritious and sufficient food has been on the rise since 2015, well before the COVID-19 pandemic exacerbated existing global food security challenges and increased the prevalence of undernourishment (PoU) by +1.5% to ~9.9% in 2020 (1,2). Food insecurity has increased in all global regions since 2014, except in North America and Europe, and the world is considerably off track to achieve the UN’s global goal of Zero Hunger by 2030 (3). Climate change is a major driver of the observed rise in global hunger (4), with both gradual climatic change and punctuated shocks working to multiply threats and disrupt the food system at various levels, most notably food production and availability, but also in terms of access, quality, utilization, and stability (5). Climate adaptation solutions are needed to lessen risk in the global agricultural production system, particularly in regions with high populations of undernourished peoples (6).

In the face of changing climate conditions and increases in the frequency and intensity of extremes, scaling up the production of NUS is an attractive strategy to build global food system resilience, particularly in the tropics and sub-tropics. Studies suggest that the agricultural adaptations made between 1981-2010 were insufficient to offset the negative impacts of climate change on common staple yields and production losses (e.g., wheat, rice, soybean, and maize), particularly at lower latitudes (7). While the geography of these major crops may expand to higher-latitudes under various climate change scenarios over the 21^st^ century (8), production of major crops in low latitudes is projected to be consistently and significantly negatively impacted (9–11), leading to a projected decrease in food availability and dietary diversity in tropical and subtropical regions (12). Furthermore, many of the world’s current food security challenges lie within tropical latitudes, where anthropogenic warming is also projected to impact agricultural production most substantially (6,12). These regions are particularly sensitive to increases in temperature due to low historical variability, with the time of emergence (TOE) of unprecedented temperature and precipitation regimes arriving earlier in terrestrial tropical biomes, compared to higher latitude regions, and potentially crossing a signal-to-noise threshold of 2 as early as the 2020s (13–15). Tropical agricultural production has already been adversely affected by extreme weather events and increased climatic variability, and yields will likely worsen with increased temperature stress and rainfall variability (16). Given these realized and projected impacts on staple crops, scaling up production of NUS has been suggested as an important adaptation measure in food-insecure regions. Expansion of NUS agriculture has the potential to enhance the climate resilience of food systems while simultaneously improving other negative environmental and social characteristics of current global food systems, including improving global crop and varietal diversity, increasing water efficiency, reducing excessive use of agrochemicals, increasing more locally produced food, and implementing practices that protect and conserve traditional knowledge and biodiversity (6).

A traditional crop originating in Oceania, where it is an important and widely used staple, breadfruit is a tropical NUS in other regions where it has a promising role in alleviating present and future food insecurity and improving the overall sustainability and resiliency of tropical and sub-tropical food systems (17,18). Breadfruit offers practical advantages in low-latitude regions as a nutritious “superfood” grown from a long-lived tree that does not need annual replanting, requires lower energy input to cultivate (especially when cultivated in sustainable agroforestry systems), sequesters carbon as a long-lived tree, and can be used as a staple food in a variety of dishes and cultural cuisines (19). Breadfruit compares favorably, in some cases surpassing, major staples (including wheat, rice, and corn) in both yield and nutritional content (20). Breadfruit also shows great potential to reduce economic barriers to food access when implemented as part of economically sustainable development strategies in food-insecure regions of the world (21). As such, breadfruit has been presented as a potential solution to the world hunger crisis and labeled a “superfood of the future” by popular media, although its potential implementation as a practical adaptation to food insecurity under a changing climate has not been thoroughly explored. That said, its potential for more intensive cultivation over its broad current range and expansion to new regions shows promise as a climate adaptation strategy for sustainable food systems. Because the most commonly grown breadfruit varieties are seedless and are clonally propagated via root cuttings, grafting, air layering, or micropropagation methods, they have little to no likelihood of becoming invasive when introduced to new regions. In fact, over the last more than ~200 years seedless varieties have been introduced to many tropical regions with no recorded invasive behavior.

Breadfruit is a domesticated crop with a rich global history, connecting various cultures and regions. Its wild, seeded ancestor is believed to have originated in New Guinea, and the seedless breadfruit was domesticated in the islands of Oceania, where hundreds of cultivars exist (22). Breadfruit played a major role as a key staple food in the human colonization of these islands. By about 3,000 years ago humans had reached Samoa and few-seeded and seedless varieties began to appear. When Hawaii was one of the last Pacific islands to be colonized by humans (between 1400 and 1700 years ago), seedless breadfruit was brought by Polynesian voyagers to islands, where it is known as “ulu” (23). Breadfruit continues to play a large part of cultural and spiritual life throughout Oceania. Europeans encountered breadfruit in Oceania in the 1500s (24). By the 18^th^ century it had been introduced to Central and South America, the Caribbean, tropical Asia, and other tropical regions, and it remains especially important in the Caribbean today (25). Given the success of breadfruit cultivation in various tropical and subtropical regions around the world, farmers in sub-Saharan Africa are increasingly looking to breadfruit as a potential crop for alleviating local food insecurity concerns (26). Despite its large promise, breadfruit remains more of a subsistence crop than commercial commodity in most regions (e.g., parts of West Africa, Southeast Asia, Latin America, and India) with major production areas limited to the Pacific and Caribbean Islands (Ragone, 2018). However, interest in commercial uses is growing (27).

The potential expansion of breadfruit cultivation as an adaptation strategy is constrained by present and future climatic uncertainties. A newly planted breadfruit tree takes several years before it begins fruiting and may live and produce fruit for over 50 years—if grown within a suitable environment. While global increases of temperature in some regions may intuitively seem to allow for the expansion of breadfruit into areas of higher latitudes, uncertainties around extreme weather, increased variability, and rising temperatures within the current cultivated extent may concurrently limit its future suitability in its current range. However, breadfruit yields are still expected to be more resilient than important global staples like rice to certain climate change stressors. For example, after a breadfruit tree is established, it may withstand droughts of 3-4 months (28). The extent of breadfruit’s resilience must be evaluated amidst an uncertain climactic future.

To examine the adaptation potential of breadfruit cultivation, we evaluate current and projected breadfruit niche suitability and assess the potential for range expansion and/or contraction using species distribution modeling (SDM) and bioclimactic data. Given our knowledge of suitable climate conditions for breadfruit cultivation, and our understanding that many of these underlying factors are undergoing anthropogenically-driven changes, we use a weighted ensemble of six species distribution models (SDMs) created using 7 selected bioclimactic variables (from BIOCLIM) to assess changes in future (2061-2080) breadfruit suitability range using downscaled simulation data from 8 different General Circulation Models (GCMs) from phase 6 of the Coupled Model Intercomparison Project (CMIP6) (29). SDMs that rely on bioclimactic variables over the full extent of a species range facilitate a realization of the fundamental niche rather than a realized niche. Consideration of seasonality and variability is useful in determining the fundamental niche of plant species, particularly when an SDM is applied to future climatic conditions. This approach allows us to assess the potential range of breadfruit in our current climate where it has not yet been introduced, as well as extrapolate breadfruit cultivation suitability in future climates. This ensemble SDM approach in conjunction with ensemble GCM projections allows us to evaluate the inherent inter-model differences/uncertainties in both steps of the process. Gaining an understanding of the SDM and climatic uncertainties that constrain breadfruit’s potential global range can facilitate policy design intended to mobilize new adaptation strategies for more resilient low-latitude food systems.

## 2 Materials and Methods

### 2.1 Species Distribution Modeling (SDM)

To determine current and future breadfruit niche suitability, we used observations and bioclimactic data to constrain 6 unique SDMs: Generalized Linear Model (GLM), Generalized Additive Model (GAM), Multiple Adaptive Regression Splines (MARS), Random Forest (RF), Generalized Boosted Model (GBM), and Maximum Entropy (MAXENT) using the *Biomodhub/Biomod2* software version 3.4.13 (2018/2021). A weighted-mean ensemble of SDMs was used to determine current breadfruit niche suitability. The ensemble approach to species distribution modeling has gained considerable popularity and is useful for examining which bioclimatic variables determine the fundamental niche of breadfruit, allowing us to project this model to future scenarios and evaluate the changed distribution of global breadfruit suitability (31).

To constrain the current distribution of breadfruit and provide training data for our SDMs, we use Global Biodiversity Information Facility (GBIF) historical breadfruit (*Artocarpus altilis*) occurrence point data (32–35). GBIF presence points are prone to sampling bias because it allows citizen science contributions to the database. Citizen scientists are prone to misidentify *A. altilis*, particularly by confusing the plant with its close relative, seeded breadnut (*A. camansi*). To limit this bias, we downloaded only those data points that were identified as part of herbarium specimens (“Preserved specimen”) or that included picture documentation. We also filtered GBIF presence points using a number of screens. First, presence data that appeared under alternative names (taxonomic synonyms), e.g., *A. communis, A. incisa,* and *A. incisus*, were included. However, data of two close relatives (i.e., *A. camansi* and *A. mariannensis*) were not included (36). Next, we ran the GBIF points using default settings of the CoordinateCleaner, an R-package tailored for cleaning occurrence records from biological collection databases (37). This package flags geographical coordinates that are overly imprecise or have common problems (e.g., points assigned to country centroids due to automated geo-referencing, plants belonging to botanical gardens or museums, etc.). We also added datapoints from a yield study in Jamaica (Trees that Feed Foundation, 2020, unpublished data). Finally, we discarded duplicate points within common grid cells to limit spatial bias. Ultimately, over half of the breadfruit presence points downloaded from GBIF were removed. The remaining 431 presence points are used for the SDM ensemble training and prediction.

To determine the climatic variables that best predict the occurrence of current breadfruit distribution, we used the dataset of bioclimatic variables (BIOCLIM) for 1970-2000 from WorldClim Version 2.1, a database of high-resolution global climate data (39). To prevent overfitting of the final SDM, we examined the full BIOCLIM dataset of 19 variables and selected a subset of variables based on their predictive performance in individual and paired runs. We selected variables largely following the methodology detailed in Moat et al., 2017 using GLM and GAM, two of the six SDMs used for the final ensemble. Boxplots, preliminary map projections, and evaluative metrics including sensitivity, specificity, True Skills Statistics (TSS), relative operating characteristic curve (ROC), and Cohen’s Kappa statistic (KAPPA) were used to determine which combination of variables were best for model fit, i.e., which combination of bioclimatic variables best predicted the observed range of breadfruit. After evaluating variable performance, we tested resulting variable combinations for collinearity and multi-collinearity using a threshold of R^2^=0.8 and reduced the selection to 7 variables (Table 1). Annual mean temperature (BIO7) and annual precipitation (BIO12) were among the highest performing variables in the preliminary model runs using GLM and GAM. While environmental characteristics such as solar radiation and soil quality also impact breadfruit growth, they are not used to constrain breadfruit suitability in this study.

The optimal combination of variables to characterize niche suitability differs based on the spatial domain. A preliminary version of the model covered the full extent of global BIOCLIM data at 10 arcminute resolution (~100 km). However, because over 97% of our preliminary weighted model results were constrained within the latitudes of 45° N to 45° S, the final analysis presented in this study was performed within these lesser latitudinal bounds. Despite this change, our analysis covers almost the entire extent of the current breadfruit range, which is ideal for accurately capturing the fundamental niche of the species (41). A lesser geographical extent also increases the reliability of pseudoabsence data. We use a presence-only SDM that relies on pseudoabsence (background) points to model suitability because no reliable source of true absence data was available. The TSS scores of the three sets of pseudoabsence points we generated vary by 0.05. Although we latitudinally limited the geographic extent of the SDM in favor of generating more accurate pseudoabsence points, it is still difficult to produce useful pseudoabsence data at this large extent, as evidenced by the concentration of pseudoabsence points in grid-cells of less-suitable categories (Supplementary Figure 1).

We ran the 7 selected BIOCLIM variables and 431 filtered data points using the Biomod2 R package version 3.4.14 using six independent SDM methods (GLM, GAM, MARS, RF, GBM, and MAXENT.Phillips). We followed established protocols for model evaluation, execution, replication, and generation of three sets of pseudo-absence data (42). Presence and pseudoabsence data were split randomly with a ratio of 70:30 to build training and testing data separately over ten runs. The same metrics of sensitivity, specificity, TSS, ROC, and KAPPA were used to evaluate model outputs.

Resultant maps and statistics from model outputs were used to evaluate the performance of the SDMs. Output statistics were reviewed to ensure a lack of difference between model runs, which could indicate problematic outcomes, such as overfitting. After evaluation, the final SDM configuration was used to produce our suitability predictions from ten runs and three replicates. The TSS threshold of 0.7 was used to remove poorly performing runs from all five models before producing a mean ensemble prediction weighted by TSS score. Our weighted ensemble of SDMs demonstrates robust present-range predictive power, characterizing the distribution of breadfruit presences indicated by GBIF data while also predicting broader distribution within Africa and other previously unexamined regions with high sensitivity (0.96) and reasonable accuracy (specificity = 0.83). The included SDMs in this study meet established criteria (TSS = 0.79, KAPPA = 0.81, ROC = 0.93).

### 2.2 General Circulation Model (GCM) Data

Projected climate data was then used to determine niche expansion/contraction under future climate scenarios SSPs 2 and 5 (43,44). We modeled the future projections over the time period of 2061-2080 using the Biomod2 package Ensemble Forecasting. The projection was performed with the same seven selected BIOCLIM variables from the GCM data under two shared socioeconomic pathway scenarios: SSP2-4.5 (stabilization GHG emission scenario) and SSP5-8.5 (high GHG emission scenario) forcing scenarios from 8 different GCMs. The GCMs from CMIP6 were used based on availability in WorldClim 2.1 and include BCC-CSM2-MR, CNRM-CM6-1, CNRM-ESM2-1, CanESM5, IPSL-CM6A-LR, MIROC-ES2L, MIROC6, and MRI-ESM2-0.

### 2.3 Analysis and Mapping

The SDM outputs from the current (Figure 1) and the two future scenarios were processed and categorized into three: Good, Fair, and Unsuitable. Niche classes were chosen based on natural inflection points and breaks between pseudoabsence and presence data (see Supplementary Figure 1). The division between “Fair” and “Good” categories was placed at 50% to approximately reflect the inflection point of the data. The Good niche classifications assesses breadfruit suitability based on yield quality and consistency, marking locations that are conducive to the establishment of new breadfruit trees. Meanwhile, the Fair range is more liberal and represents climactic conditions that established trees will likely withstand, ergo persistence is assumed likely.

**Figure 1.**
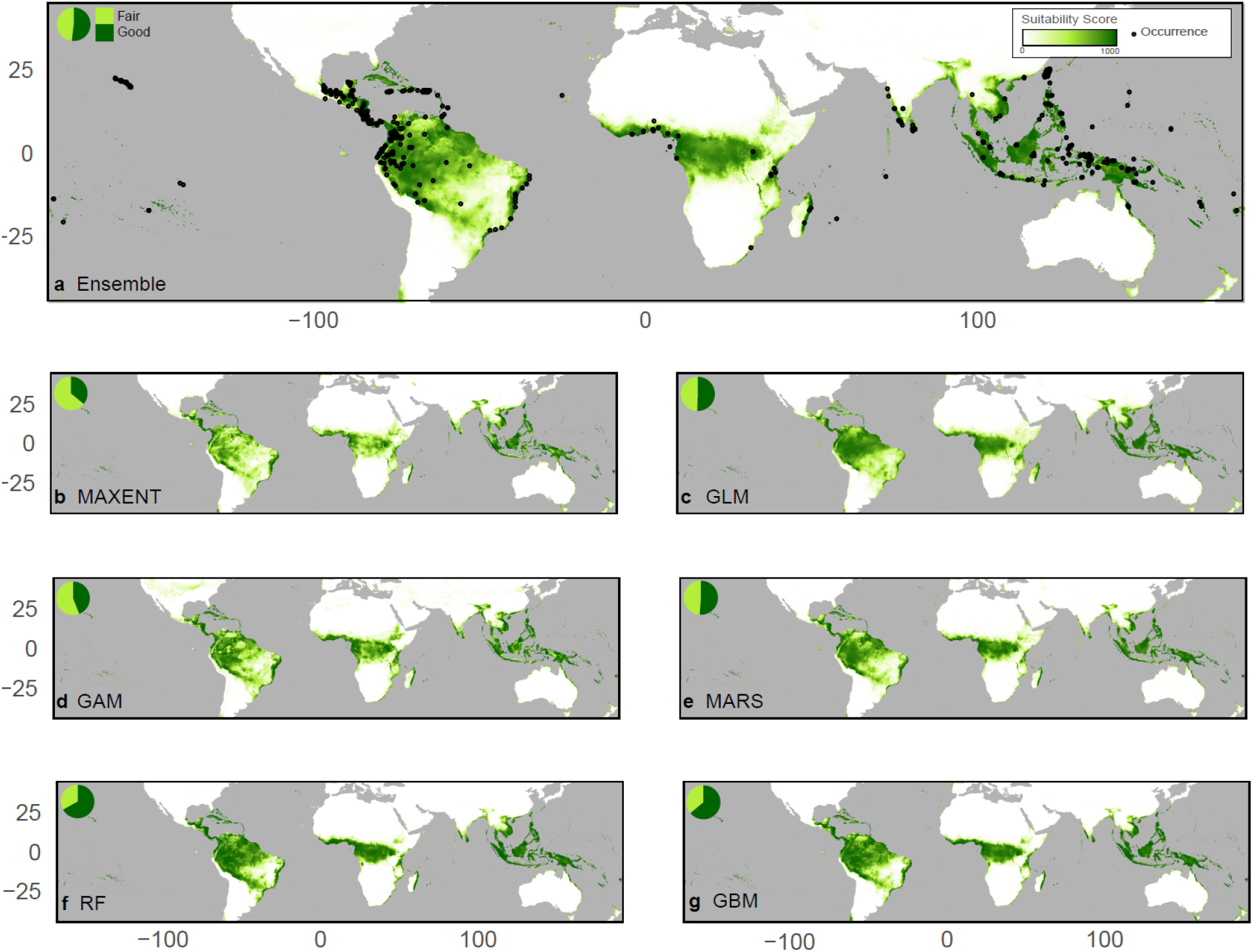
Range of current (1970-2000) breadfruit suitability in the global tropics and subtropics (45° N to 45° S) as predicted from bioclimactic variables from 10 arcminute resolution WorldClim v2.1 data and filtered GBIF occurrence points. Climatic suitability of breadfruit cultivation is indicated by green shading ranging from 0 (fair) to 1000 (good). (*A*) The ensemble mean-weighted SDM and 6 different individual SDMs including (*B*) MAXENT, (*C*) GLM, (*D*) GAM, (*E*) MARS, (*F*) RF, and (*G*) GBM. Black dots in (*A*) are observed occurrence points from GBIF. Pie charts in upper left of each plot indicate fraction of Fair (score=250-750) to Good (score=750+) area.

## 3 Results

### 3.1 SDM Prediction of Current Breadfruit Suitability Range

Our weighted ensemble baseline SDM predicts niche suitability in all GBIF-identified locations of current breadfruit cultivation, as well as substantial swathes of the global tropics and subtropics where successful cultivation of breadfruit has not yet been confirmed. Predicted suitable areas for cultivation include large sections of Southeast Asia, Central America, northern South America, the Caribbean, and equatorial Africa (Figure 1a). Latitudinal limits of modeled suitability extend as far north as northeastern India in Asia and southern Florida in North America and stretch as far south as Chile in South America and New Zealand in the Pacific. These regions have previously been identified by the National Tropical Botanic Garden (NTBG) as suitable for breadfruit cultivation based on current climatic conditions (45). Suitable niches that extend into the higher latitudes of the tropics are largely concentrated along coastlines, in agreement with available GBIF occurrence data (Figure 1a; black dots). Notable areas without widespread occurrence data that are identified as having robust potential for breadfruit cultivation include Cuba, equatorial Africa, the southeastern coastline of Africa, and the Malay peninsula. In Africa, the lack of occurrence data in GBIF may be explained by breadfruit’s recent arrival to the continent, but recent successful introductions haven been made in Uganda, Kenya, Tanzania, and Ghana. While no GBIF records are recorded in Cuba, it is known to grow there (26,46,47).

Our SDM breadfruit suitability predictions employ seven weighted bioclimatic variables to characterize the climatic conditions within a grid cell. The environmental variables were selected based on evaluation scores on the Generalized Linear Models and Generalized Additive Models while controlling for multicollinearity and overfitting (see Methods). Contribution weights of the seven bioclimatic variables are quantified using True Skill Statistic (TSS) scores. We find that baseline breadfruit suitability is largely driven by three temperature-related bioclimatic variables (TSS: 60.9). The annual temperature range, defined as the difference between the maximum temperature of the warmest month and the minimum temperature of the coldest month is the predominant prediction contributor (Figure 4c; TSS: 40.1). Secondary contributors include the mean diurnal temperature range (Figure 4a; TSS: 9.2) and the mean temperature of the coldest month (Figure 4g; TSS: 11.5). The four remaining contributing variables – annual precipitation, and precipitation during the driest, warmest, and coldest quarters – are lesser contributors relative to the temperature variables (Figure d-g; TSS: 6.1-7.0).

To refine our suitability findings and identify the niches most suitable for cultivation, we employ statistical thresholding to subdivide the SDM suitability scores into Unsuitable, Fair, and Good categories (Supplementary Figure 1). The unsuitable niche classification marks areas where breadfruit cultivation is barely possible to impossible, whereas fair classification marks areas where breadfruit cultivation is possible but yield and quality may be negatively affected by adverse seasonal weather conditions (e.g., low rainfall or high temperatures). Good niche classification indicates where breadfruit cultivation has the potential for consistent, high quality yields that are less likely to be affected by adverse weather conditions. For the lower threshold, between unsuitable and fair, we set the cut-off at the inflection point where an increase in model suitability values and the cumulative percentage of all (including pseudoabsence and occurrence) points plateaus, which resulted in the loss of four ground control points. The division between Fair and Good categories indicates a second inflection point where model suitability values begin to substantially increase.

Individual SDMs are remarkably consistent in their suitable geographic range predictions (Figure 1b-g) but show notable differences in categorizing the level of breadfruit suitability within these ranges. Across the 6 SDMs, there is only a 3.3% difference in the total “suitable” land area, although the classification of suitable land into either Fair or Good is less consistent, with ~33.1-64.0% of the total suitable area identified as Fair depending on SDM and ~36.0-66.9% classified as Good. When considering the total modeled area of suitability, both MAXENT (35.9%, 10.0M km^2^ Good; 64.1%, 17.7M km^2^ Fair) and GAM (44.0%, 12.5M km^2^ Good; 56.0%, 15.8M km^2^ Fair) predict lesser “Good” areas in proportion to “Fair”, while RF (66.9%, 8.4M km^2^ Good; 33.1%, 17.0M km^2^ Fair) and GBM (63.9%, 16.2M km^2^ Good; 36.1%, 9.1M km^2^ Fair) predict greater “Good” suitability (Figure 1). Because MAXENT is a presence-only SDM, it is known to underestimate the probability of occurrence within areas of observed presence, while overestimating in areas without known presence (48). Several MAXENT runs (6 out of 30) were removed from the final weighted SDM ensemble due to low TSS scores, likely due to a lack of convergence. Marginal areas of Fair suitability are consistently identified by all 6 SDMs in the same locations (e.g., New Zealand, southwestern coast of Africa, Chile). Southeast Asia, Oceania, and the Pacific Islands are also consistently valued with the highest proportion of Good to Fair suitability. The 6 SDMs demonstrate consistent suitability coverage and patterns in regional Good to Fair suitability, although specific values differ.

Using the weighted-ensemble projection, we focus our narrative on comparative breadfruit suitability in three main regions: Asia and the Pacific Islands, the Caribbean and Latin America, and Africa. Compared to the averaged global proportion of area with Good (51.9%, ~14M km^2^) to Fair (48.1%, ~12M km^2^) suitability (Figure 3), breadfruit is particularly suited to climactic conditions in Southeast Asia. Breadfruit is suitable for cultivation within 80.7% of the area (~3.6M km^2^) in this region with a high proportion of Good (77.4%, ~2.8M km^2^) to Fair (22.6%, ~800k km^2^) quality, although Pacific Islands such as Hawaii also show similarly high levels of Good (87.7%, ~30k km^2^) to Fair (12.3%, ~4k km^2^) area. In Latin America and the Caribbean, island countries show high coverage of suitable area (99.2%, ~2.3M km^2^) while 51.7% (~11.7M km^2^) of the overall land area in Latin America and the Caribbean is projected to be suitable. 41.3% of these suitable areas are classified as Good and 58.7% as Fair. In Africa, 20.8% of the continent is projected to be suitable for growing breadfruit, with 45.4% of this land showing Good and 54.6% showing Fair suitability. This baseline SDM prediction gives a comprehensive picture of the range and quality of suitable breadfruit-growing areas in various low-latitude regions of the world.

### Change in Breadfruit Range due to Climate Change

To assess the effects of climate change on breadfruit suitability, we project our SDM to future (2061-1080) climate conditions using an ensemble of opportunity of 8 GCMs from CMIP6. While breadfruit is projected to largely maintain its present range, the resilience and adaptation potential is projected to decrease, with more substantial decreases in Good area projected from the high-emissions scenario (44.4%) than the stabilization scenario (32.1%) (Figure 2). We find a 4.7% decrease in weighted ensemble SDM suitable breadfruit range (~1.2M km^2^) in the stabilization scenario and a 4.5% decrease (~1.2M km^2^) under high emission scenario conditions (Figure 3). However, the total area of “winners” and “losers” in terms of overall breadfruit suitability changes largely balance on a global scale.

**Figure 2.**
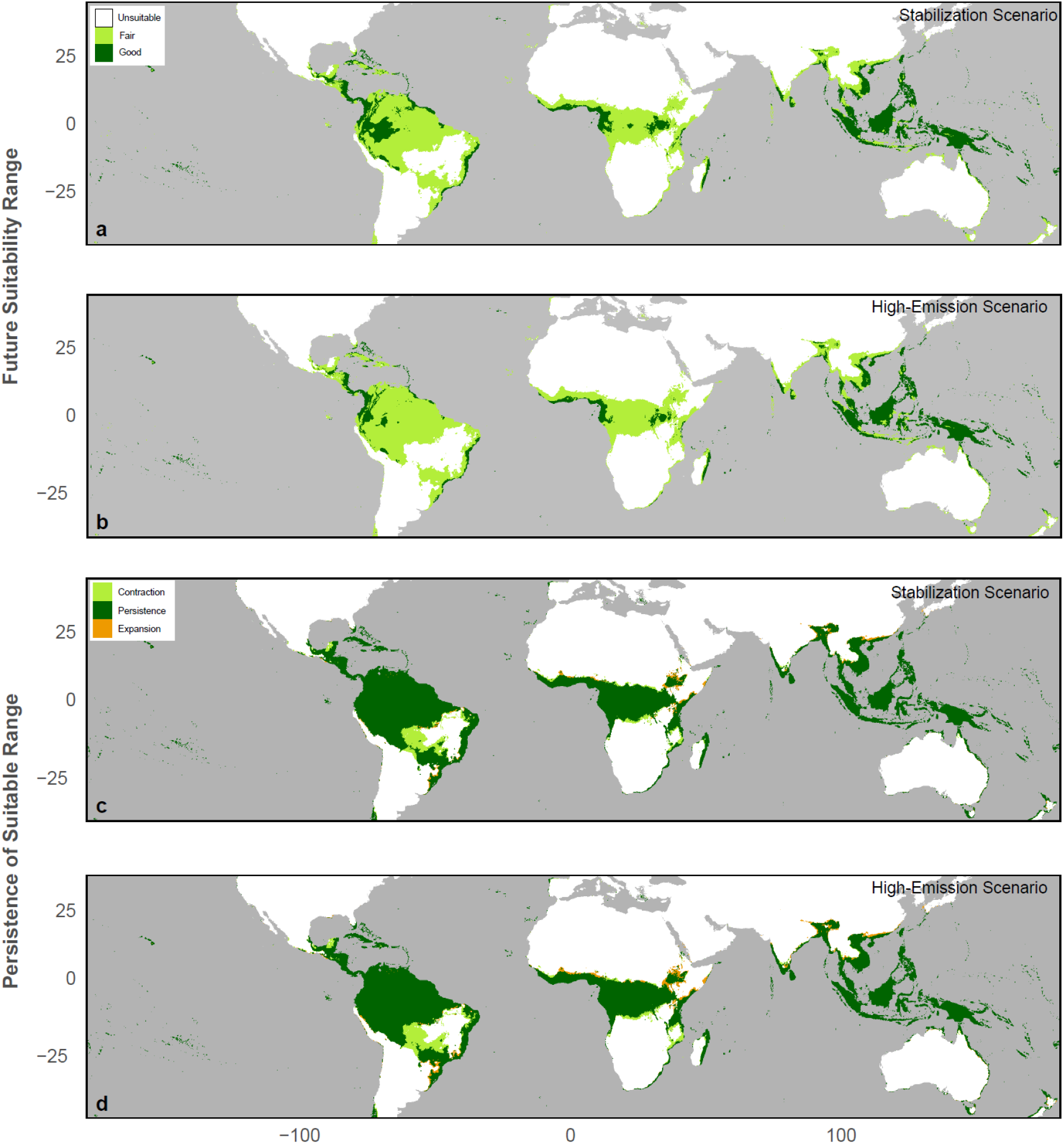
Future (2061-2080) breadfruit suitability range and change under stabilization and high-emission scenarios. (a) Future suitability (Fair: score=250-750; Good: score=750+) of weighted SDM ensemble under stabilization scenario (SSP2-4.5) and (b) high-emission scenario (SSP8-8.5) using averages from a CMIP6 ensemble of opportunity from 8 GCMs. Change in suitable range (score>250) from baseline suitability score under (c) stabilization scenario and (d) high-emission scenario.

**Figure 3.**
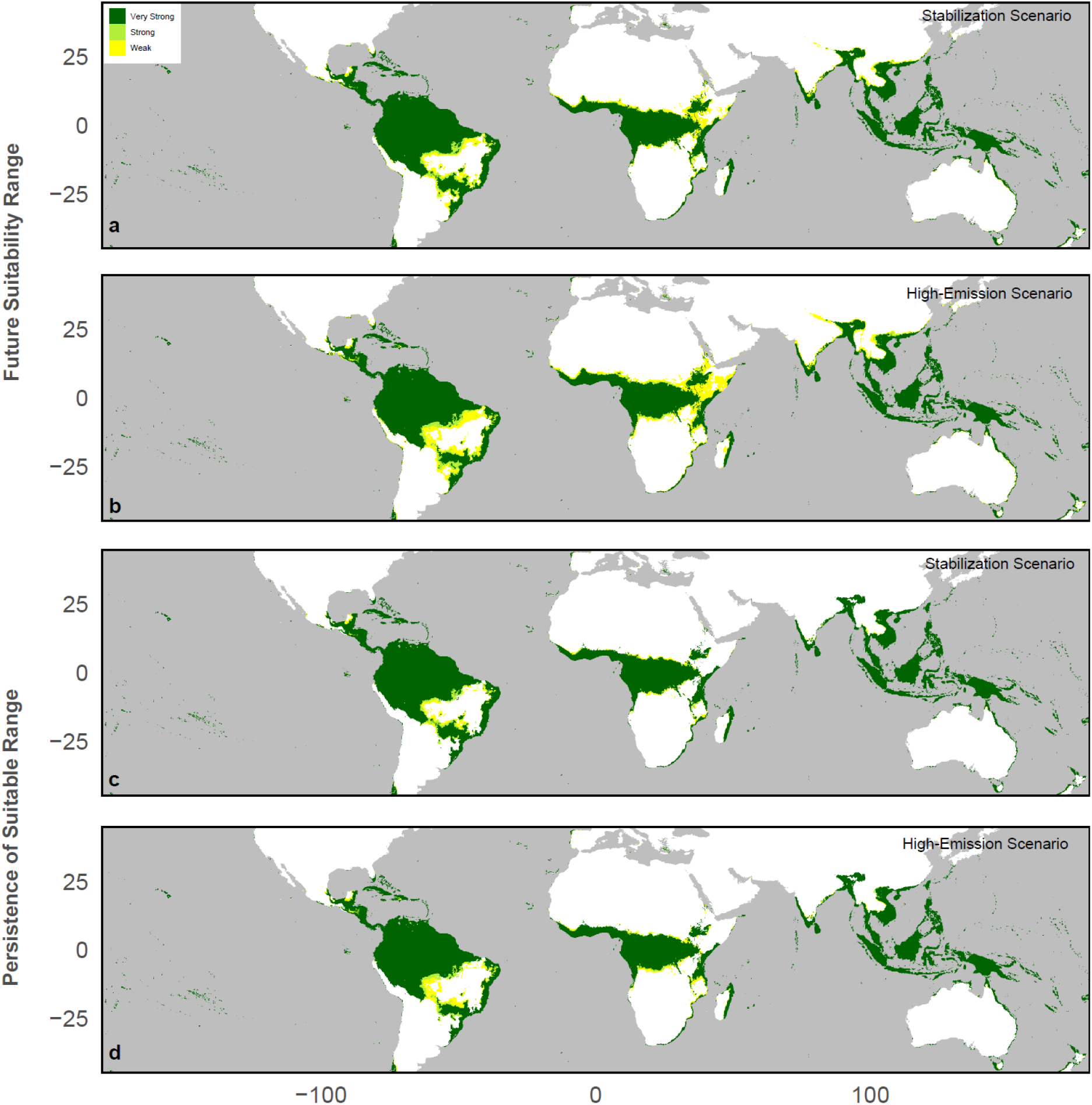
Change in suitable breadfruit area (value>250) between the ensemble baseline SDM (B) and the stabilization (S) and high-emission (H) scenarios within various regions in the global tropics and subtropics, including Asia and the Pacific Islands, Africa, and Latin America, with Weak (1-3 members), Strong (4-5 members) and Very Strong (6+ members) CMIP6 GCM average agreement.

Change in suitable breadfruit range – expansion, contraction, and persistence— from the baseline projection varies considerably across regions (Figure 2). The suitable area primarily persists and even increases ~0.7% (~23k km^2^) under the stabilization scenario and 0.8% (~30k km^2^) under the high emission scenario in Southeast Asia, although this increase in area is accompanied by a decrease in quality (2.6% and 8.7%) and reduction in total Good area. In Asia and the Pacific Islands more broadly, total suitable breadfruit area increases ~2.8% (~144k km^2^) and ~4.1% (~214k km^2^), showing areas of potential expansion in the upper and lower latitudinal limits of this region (Figure 2C, D). The majority of suitable breadfruit-growing area is projected to persist in equatorial Africa, although with lower resilience potential to future climate conditions than in Southeast Asia (1.5-3.4% reduction in total suitable area and 58.2-70.0% reduction in Good area depending on emissions scenario). In both the stabilization and high-emission scenarios, the baseline suitability range within equatorial Africa expands northward while the southern bounds of breadfruit suitability are projected to contract. However, Latin America and the Caribbean are projected to experience the largest magnitude range reduction on both the stabilization (10.1%) and high emission pathways (11.5%). These range reductions appear to be concentrated in specific areas, particularly in the Yucatan peninsula and southwestern Brazil. While Caribbean islands experience minimal contraction of the overall suitable breadfruit-growing area (1.4-1.6% reduction) under either emission pathway, the quality of these conditions may decrease significantly (50.6-72.5% reduction in Good area). Despite regional variation, a large portion of global suitable breadfruit range is projected to persist in both scenarios (particularly in Africa and Asia and the Pacific Islands), suggesting that (1), established trees will likely withstand future climatic conditions in most of the tropics and subtropics and (2) there is room for expanded cultivation of breadfruit across a broad section of the global tropics and subtropics in Africa. However, a greater decrease in quality but not necessarily a greater decrease in total area of suitability is projected under the high emissions scenario in all regions.

Projected expansion of breadfruit suitability is driven by increases in seasonal temperature and precipitation. Increased precipitation during the coldest quarter (BIO19, Fig. 4t, u) as well as general increases in annual precipitation (BIO12, Fig. 4k,l) drive potential expansion of suitable breadfruit area at the perimeter of the baseline suitable range. Meanwhile, decreases in precipitation of the warmest quarter (BIO18) drive contraction of suitable breadfruit range in Central and South America (Fig. 4q, r). Mean Diurnal Range shows little change in Southeast Asia but increases in areas of expansion and decreases in areas of contraction and reduced breadfruit suitability (Fig. 4b, c).

**Figure 4.**
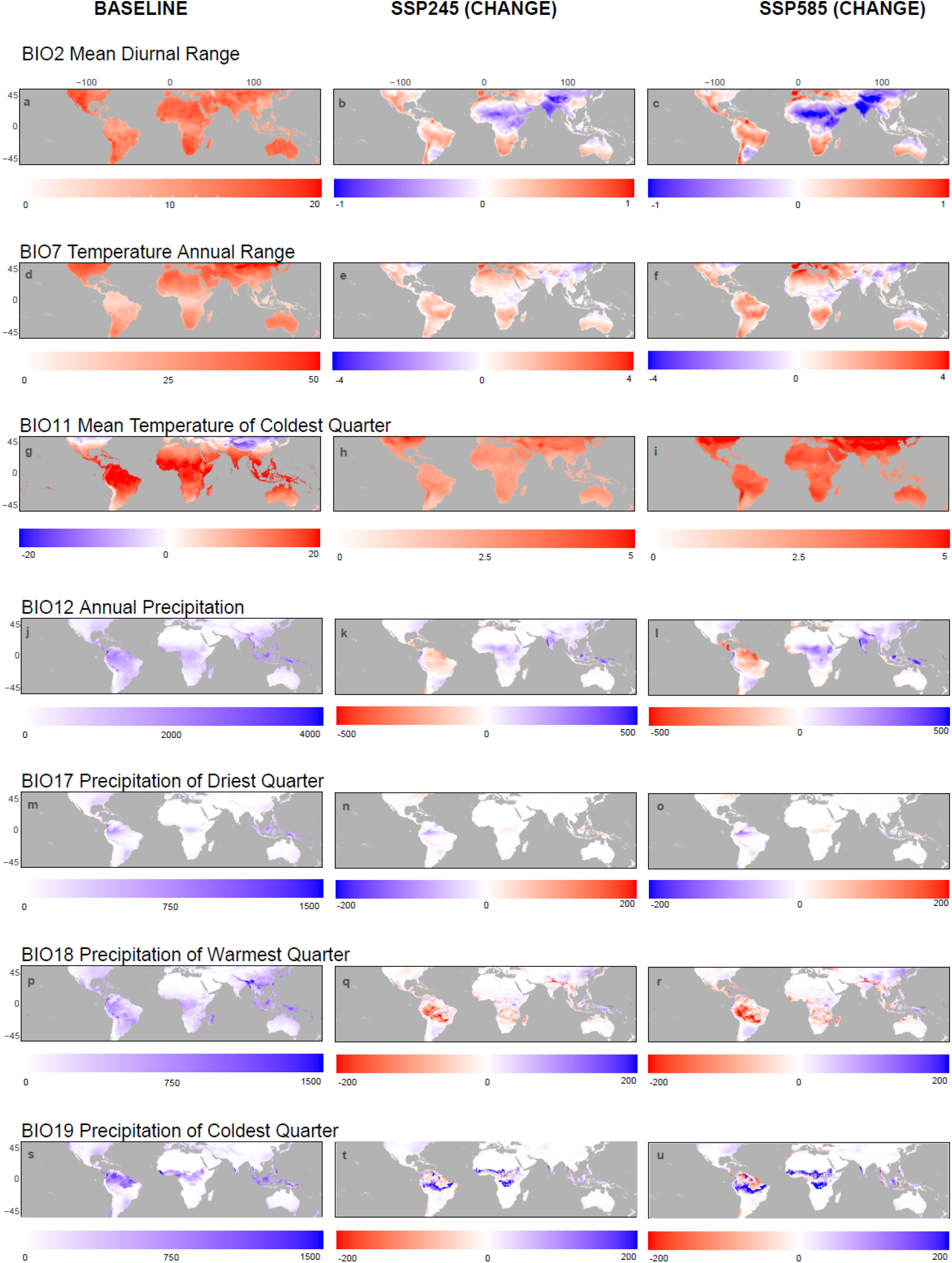
Difference Plots of Top BIOCLIM Contributors to SDM. Columns show the baseline (1970-2000) bioclimatic variables (left) and the absolute difference of the specific bioclimatic variables from 2061-2080 under SSP2-4.5 (middle) and SSP5-8.5 (right) scenarios of the CMIP6 ensemble of opportunity downscaled to 10 arcminute resolution from WorldClim v2.1.

To characterize the structural uncertainty of our breadfruit projections, we use our CMIP6 ensemble of opportunity and define projection confidence according to ensemble member agreement (i.e., 4-5 members agree – Strong, 6+ members agree = Very Strong). Projections of future suitability show Very Strong GCM agreement over the majority of the persistent range of breadfruit suitability under the stabilization scenario (92.8%) and high-emission scenario (90.5%), with a larger area of weak GCM agreement under the high-emission scenario. Weak SDM agreement (<4 members agree) only characters 5.1% of the persisting suitable range under the high-emission scenario and 3.6% of the suitable range under the stabilization scenario. Strong and Weak agreement is concentrated at the outer fringes of breadfruit distribution and most significantly in central South America in the Mato Grosso Plateau and eastern Africa in the Ethiopian highlands (Figure 3). This finding suggests higher levels of uncertainty in areas of higher elevation, although breadfruit suitability is projected to persist in the Andes in South America. Areas of potential expansion also overlap with the areas of greatest uncertainty. In future scenarios, South Asia and South America exhibit the greatest variability in area among GCMs (Figure 4). However, the majority of future global breadfruit cultivation in both scenarios demonstrates strong (4-5) to Very Strong (6+) agreement among the 8 GCMs in both scenarios.

## Discussion

Our SDM prediction captures known locations of breadfruit while also suggesting potential expansion for the crop into food-insecure regions, most notably within Sub-Saharan Africa. While previous studies have attempted to constrain global and regional distributions of breadfruit suitability (18,45), direct comparisons are challenging due to differences in scoring systems. However, the geographic range of predictions from other studies are in good agreement with our ensemble SDM predictions. For example, Lucas and Ragone (2012) used Geographic Information Systems (GIS) to produce a map of global breadfruit suitability using WorldClim v1.4 data and ecologically determined ranges for mean temperature and rainfall (Ragone, 2006). While this study did not assess the impacts of climate change, their results on the current distribution of suitable cultivation range is in good agreement with our SDM-based findings. Mausio et al. (2020) further used fuzzy-set models to extrapolate breadfruit suitability from environmental parameters derived in Hawaii. Their work offers a useful comparison of an alternative approach to modeling breadfruit suitability by incorporating FAO EcoCrop parameters and climate data from CMIP5, but it does not address the impact of seasonal temperature and precipitation data on potential breadfruit distribution under future scenarios.

Our SDM indicates that the differences between the high-emissions scenario and stabilization scenario in terms of total breadfruit suitability area is minimal, however the high-emission scenario is predicted to lead to more substantial reductions in suitability scores in most regions. These findings are particularly important for tropical and subtropical regions (low to mid latitude regions) where impacts on the production of conventional staple crops such as maize, wheat, soybean and rice are projected to be more negative than higher latitudes (49). Expanded cultivation of breadfruit in such regions may contribute to global food and economic security, as a locally produced staple food that can lead to greater independence from food imports.

Breadfruit shows the most promising resilience to future climactic conditions within its native extent in Asia, largely persisting in Southeast Asia and even showing potential areas of expansion within India and China. Culturally, breadfruit is also a significant traditional crop in many Pacific Islands including Hawaii, where it has been noted for its promising role as part of local biocultural restoration efforts (Langston and Lincoln, 2018). Breadfruit can play an important role in local low-latitude food systems, alleviating the yield losses of major crops such as rice, while promoting sustainable agroforestry systems. However, the persistence of suitable conditions for growing breadfruit are most at risk in specific regions within Latin America, namely a section of the Yucatan peninsula and southwestern Brazil in the Mosso Plateau. These areas experience decreases in annual precipitation, with significant decreases in precipitation during the warmest quarter of the year. These areas also coincide with weak agreement among the 8 GCMs, with less than half agreeing on the magnitude of precipitation reduction.

Our SDM predictions also signal opportunities to expand cultivation in Central Africa, building on nascent cultivation of breadfruit which currently exists in parts of West and East Africa (26). Given the present state of food insecurity in Africa, the widespread cultivation of breadfruit offers a path towards increased food security and independence in the present and more distant future for several Sub-Saharan countries. As a promising NUS, breadfruit could fill immediate and future needs to increase food security in many tropical regions, especially as crops such as maize and rice become less suited to warming conditions (11,50). Breadfruit demonstrates resilience to uncertain climactic conditions in many tropical food-insecure regions. This adaptation potential is further enhanced if breadfruit trees are well-established and grown within sustainable agroforestry systems. Agroforestry systems can serve as both adaptation and mitigation, contributing to food productivity while enhancing restoration by combining traditional growing methods used by Pacific Islanders for centuries with modern horticultural techniques (6,51).

Our study uses species distribution modeling to examine potential areas for breadfruit cultivation over the next several decades under two emission pathways. Our model prediction is limited to bioclimactic factors (seasonal temperature and precipitation variables) and does not include the potential influence of soil type, solar radiation, or elevated CO_2_ concentrations on breadfruit suitability. We are confident that the 45° N to 45° S swath covers the majority (~97%) of present and future suitable breadfruit range under these scenarios, capturing the fundamental niche and identifying new areas where breadfruit can be introduced or expanded globally. While additional presence observations may improve the performance of our model, most studies agree that sample size effects usually become less critical above 50 presence points (Guisan et al., 2017). Ideally, models of habitat suitability should also be based on measurements of population fitness at each geographic location. However, this would prevent the use of reliable data available in natural history collections. Although data points from GBIF were cleaned and filtered, absence data and rigorous groundtruthing would further improve the performance of the individual models that made up the final SDM ensemble. When projecting to future conditions, we were limited to using realizations from relatively few (8) CMIP6 GCMs from WorldClim v.2.1 which likely limits the range of internal variability and structural uncertainty considered in our projections.

Significant portions of sub-Saharan Africa show room for expansion of breadfruit cultivation over the next several decades, especially as rapid population growth is projected to intensify food insecurity and undernourishment across the continent (52). Closing the yield gap via sustainable intensification of agriculture which utilizes NUS such as breadfruit may be critical in preventing future catastrophic circumstances. Prioritizing a rich diversity of NUS over staple crops such as wheat, maize, and rice will lead to healthier diets and improved nutrition and contribute to sustainable development (53). Similar studies have examined the potential distribution of NUS under climate change, including 4 neglected and underutilized fruit species (NUFS) in Sri Lanka and a yam species in Benin (54,55), but few have examined the viability of expanding cultivation of NUS across different continents at a global scale. As a long-lived crop that can be grown and used locally, breadfruit offers an opportunity for various low-latitude communities around the world to improve the resilience of their local food systems. By building agroforestry systems around breadfruit, low-latitude regions can avoid the pitfalls of strictly monoculture systems while building sustainable food systems with increased resilience to climate change.

## 4.4 Conclusion

Our model of breadfruit niche suitability has the potential to provide projections useful for both short-term decision making and long-term adaptation planning. Because breadfruit cultivation has not yet expanded into its full suitable range, this crop has the potential to alleviate present and future food security burdens, particularly in food-insecure countries in equatorial Africa. Although breadfruit growers may need to contend with a certain level of reduced quality of yields in the future, suitable climatic conditions for breadfruit cultivation exists across a broad range of the tropics and subtropics and persist for the next several decades regardless of emissions pathway, indicating that it is a reliable and promising NUS to implement as a climate adaptation strategy. Depending on scale of production, breadfruit itself may even play a role in contributing to climate change mitigation due to its significant carbon sequestration benefits.

## Acknowledgments

Research presented in this paper was supported by Summer Undergraduate Research and Conference Travel Grants from Northwestern University’s Office of the Provost to LY. The authors declare that there is no conflict of interest.

---

**Supplementary Figure 1.**
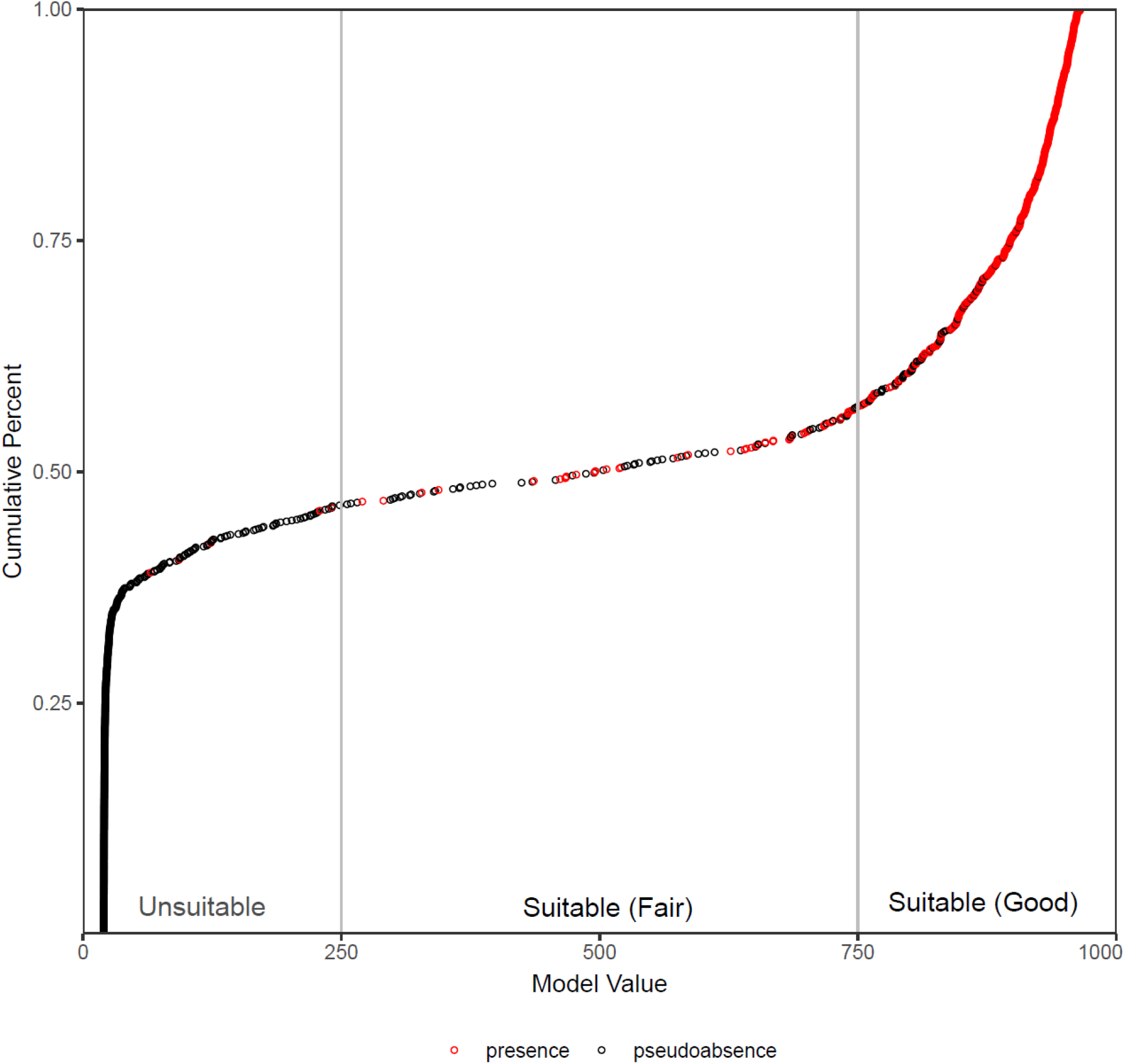
Threshold metrics for breadfruit suitability used for SDM classification. Categories are demarcated in relation to trends between cumulative percent of total number of points (presence and pseudoabsence) and model suitability value (0-1000). The Fair (250-750) and Good (750-1000) suitability categories are marked by greater prevalence of presence points, with 4 occurrence points categorized as unsuitable.

**Supplementary Table 1.** Relative importance (TSS) of 7 selected BIOCLIM variables within and across 6 individual SDMs in baseline (1970-2000) scenario.

|  | Bioclimactic Variable | MAXENT | GLM | GAM | MARS | RF | GBM | Average |
| --- | --- | --- | --- | --- | --- | --- | --- | --- |
| BIO2 | Mean Diurnal Range | 0.201 | 0.046 | 0.123 | 0.081 | 0.063 | 0.042 | 0.093 |
| BIO7 | Temperature Annual Range (BIO5-BIO6) | 0.312 | 0.543 | 0.403 | 0.381 | 0.312 | 0.454 | 0.401 |
| BIO11 | Mean Temperature of Coldest<br>Quarter | 0.194 | 0.052 | 0.191 | 0.124 | 0.098 | 0.033 | 0.115 |
| BIO12 | Annual Precipitation | 0.232 | 0.115 | 0.216 | 0.130 | 0.057 | 0.029 | 0.130 |
| BIO17 | Precipitation of Driest Quarter | 0.121 | 0.027 | 0.106 | 0.055 | 0.031 | 0.025 | 0.061 |
| BIO18 | Precipitation of Warmest Quarter | 0.150 | 0.063 | 0.107 | 0.026 | 0.038 | 0.016 | 0.067 |
| BIO19 | Precipitation of Coldest Quarter | 0.185 | 0.012 | 0.104 | 0.069 | 0.034 | 0.016 | 0.070 |

**Supplementary Table 2.**
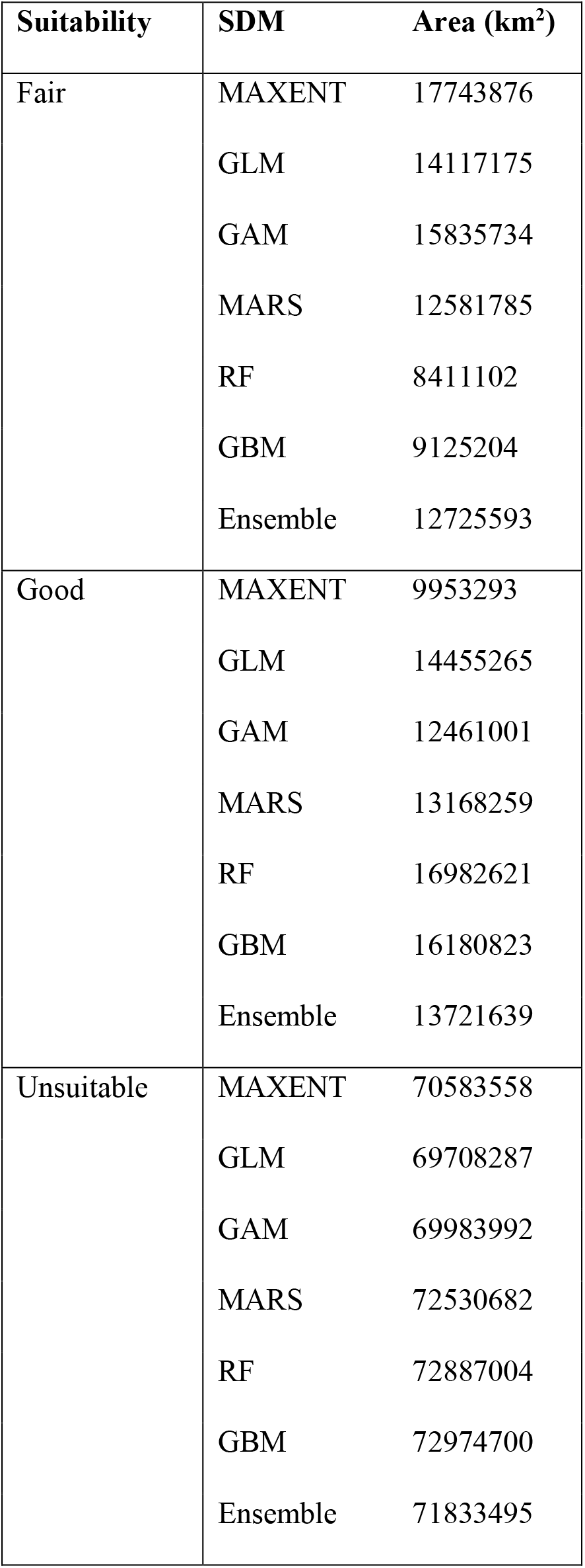
SDM Variability of Breadfruit Suitability across 6 individual and ensemble SDMs. Suitability is categorized over 98280727 km^2^ of total land area constrained within the latitudes of 45° N to 45° S.

| Suitability | SDM | Area (km<sup>2</sup>) |
| --- | --- | --- |
| Fair | MAXENT | 17743876 |
|  | GLM | 14117175 |
|  | GAM | 15835734 |
|  | MARS | 12581785 |
|  | RF | 8411102 |
|  | GBM | 9125204 |
|  | Ensemble | 12725593 |
| Good | MAXENT | 9953293 |
|  | GLM | 14455265 |
|  | GAM | 12461001 |
|  | MARS | 13168259 |
|  | RF | 16982621 |
|  | GBM | 16180823 |
|  | Ensemble | 13721639 |
| Unsuitable | MAXENT | 70583558 |
|  | GLM | 69708287 |
|  | GAM | 69983992 |
|  | MARS | 72530682 |
|  | RF | 72887004 |
|  | GBM | 72974700 |
|  | Ensemble | 71833495 |

**Supplementary Table 3.**
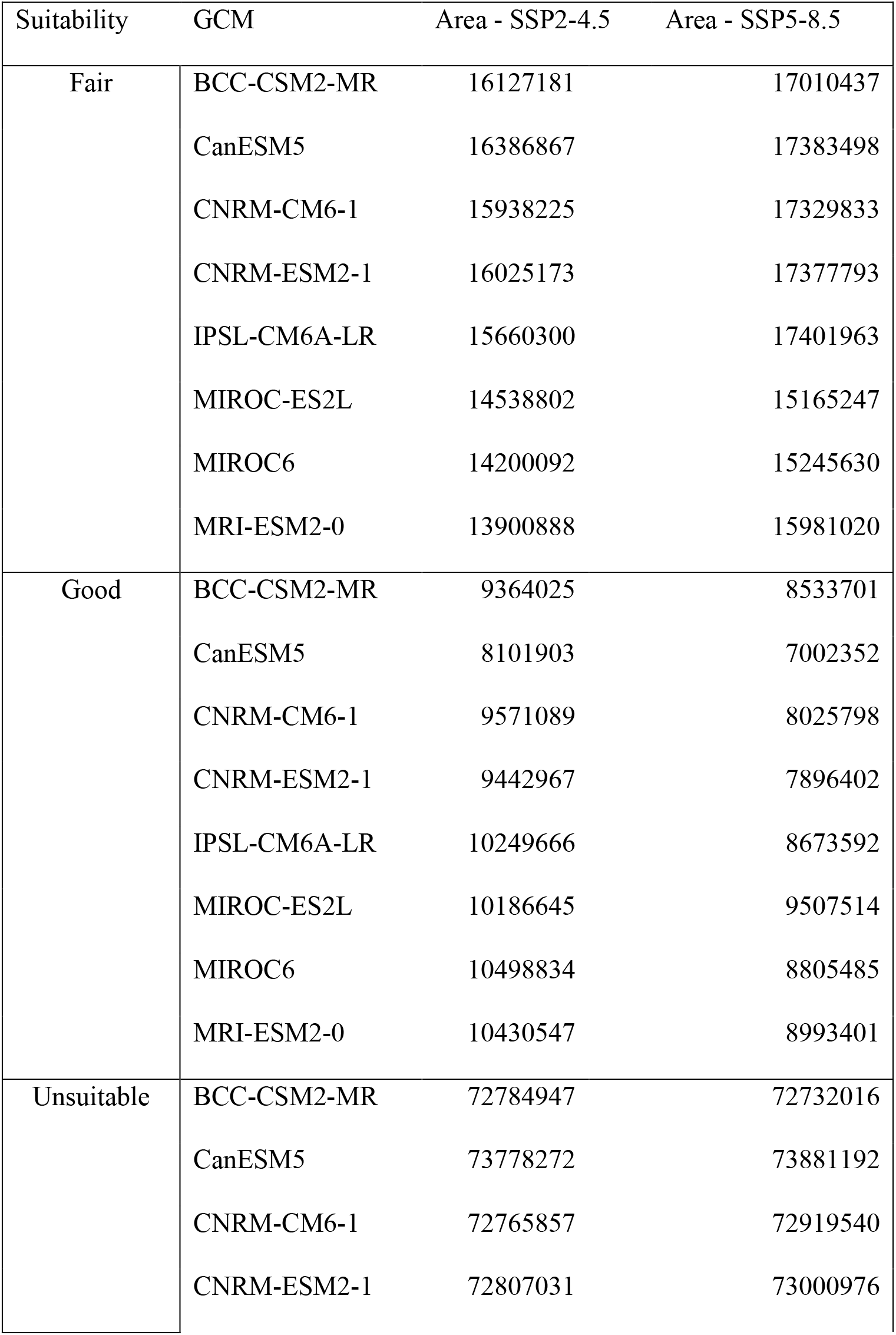
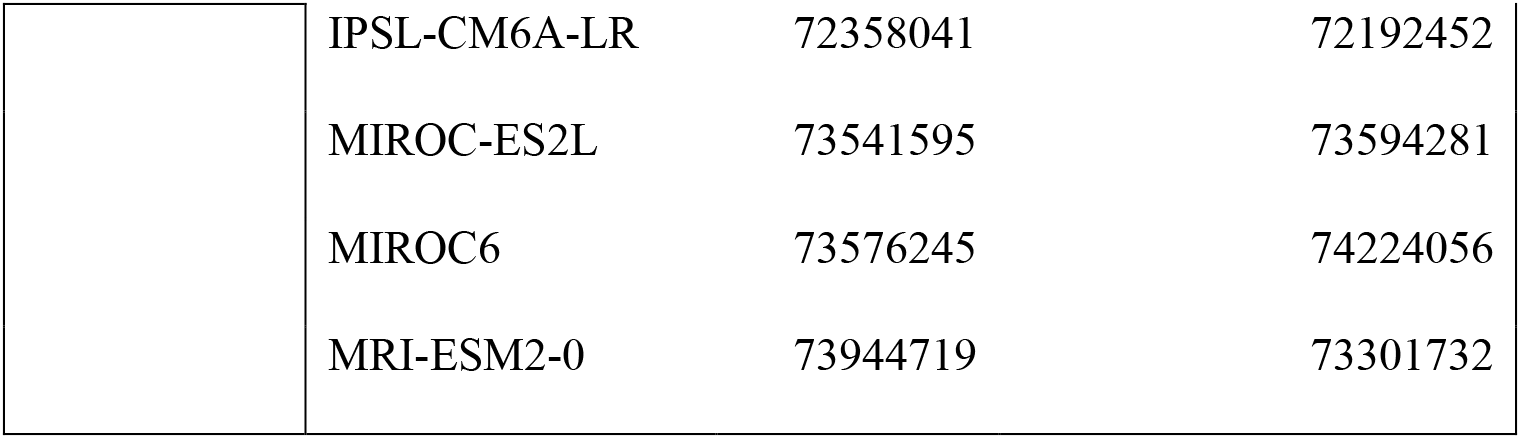
GCM Variability in Breadfruit Suitability Under Stabilization and High Emissions Scenarios of individual CMIP6 GCMs. All available downscaled GCMs at 10 arcminute resolution were accessed from WorldClim v2.1. Breadfruit suitability is evaluated within land area constrained within the global tropics and subtropics.

| Suitability | GCM | Area - SSP2-4.5 | Area - SSP5-8.5 |
| --- | --- | --- | --- |
| Fair | BCC-CSM2-MR | 16127181 | 17010437 |
|  | CanESM5 | 16386867 | 17383498 |
|  | CNRM-CM6-1 | 15938225 | 17329833 |
|  | CNRM-ESM2-1 | 16025173 | 17377793 |
|  | IPSL-CM6A-LR | 15660300 | 17401963 |
|  | MIROC-ES2L | 14538802 | 15165247 |
|  | MIROC6 | 14200092 | 15245630 |
|  | MRI-ESM2-0 | 13900888 | 15981020 |
| Good | BCC-CSM2-MR | 9364025 | 8533701 |
|  | CanESM5 | 8101903 | 7002352 |
|  | CNRM-CM6-1 | 9571089 | 8025798 |
|  | CNRM-ESM2-1 | 9442967 | 7896402 |
|  | IPSL-CM6A-LR | 10249666 | 8673592 |
|  | MIROC-ES2L | 10186645 | 9507514 |
|  | MIROC6 | 10498834 | 8805485 |
|  | MRI-ESM2-0 | 10430547 | 8993401 |
| Unsuitable | BCC-CSM2-MR | 72784947 | 72732016 |
|  | CanESM5 | 73778272 | 73881192 |
|  | CNRM-CM6-1 | 72765857 | 72919540 |
|  | CNRM-ESM2-1 | 72807031 | 73000976 |
|  | IPSL-CM6A-LR | 72358041 | 72192452 |
|  | MIROC-ES2L | 73541595 | 73594281 |
|  | MIROC6 | 73576245 | 74224056 |
|  | MRI-ESM2-0 | 73944719 | 73301732 |

**Supplementary Figure 2.**
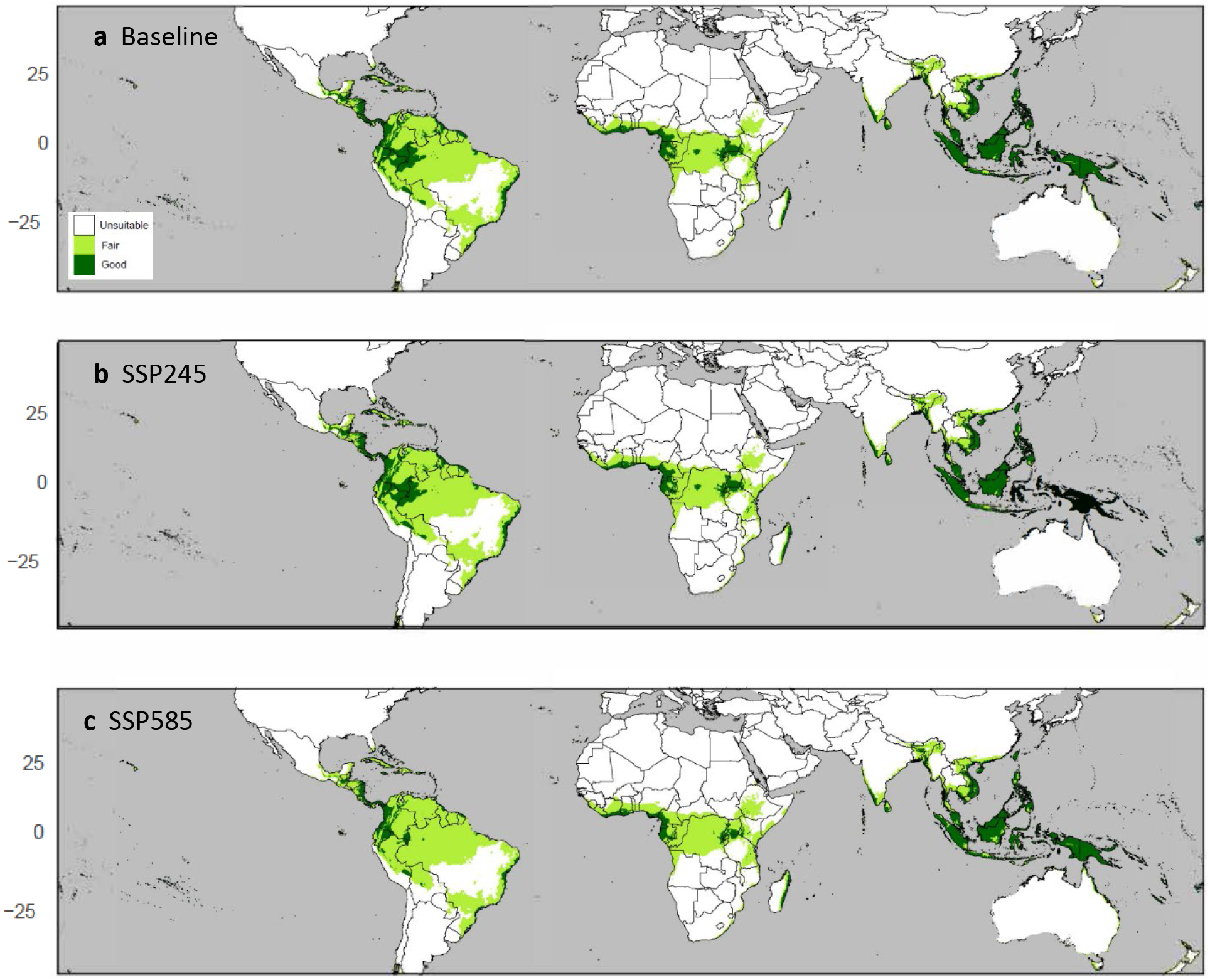
Suitable Breadfruit Range. (a) baseline (1970-2000), (b) Future Stabilization (2061-2080), and (c) Future High Emissions (2061-2080) scenarios with country outlines using average values from a weighted SDM ensemble projected using a CMIP6 GCM ensemble of opportunity under SSPs 2 and 5.

